# Lepidoptera proboscis pollens are mainly derived from nectar

**DOI:** 10.1101/2023.03.26.534265

**Authors:** Baiwei Ma, Qi Chen, Fei Lin, Guirong Wang, Bingzhong Ren

**Author notes:** **Corresponding author: Bingzhong Ren (BZR):** Jilin Provincial Key Laboratory of Animal Resource Conservation and Utilization, School of Life Sciences, Northeast Normal University, Changchun, CO130024, China., **Guirong Wang (GRW):** Shenzhen Branch, Guangdong Laboratory for Lingnan Modern Agriculture, Genome Analysis Laboratory of the Ministry of Agriculture, Agricultural Genomics Institute at Shenzhen, Chinese Academy of Agricultural Sciences, Shenzhen, CO518120, China.

## Abstract

The pollens which are rejected or ignored by the dry stigma can’t germinate since they haven’t acquired sugar solution from dry stigmas. But almost all research has ignored the participation of insects which may transport pollens from the nectar to stigmas, and that pollens may germinate on the dry stigma since they have soaked in the nectar. So, the key question is that whether insects carry pollens from nectar to the stigma.

Since the adult Lepidoptera are important to the plant pollination and there is a general consensus that pollens are mainly deposited on the proboscis of adult Lepidoptera, and pollens are ubiquitous in the nectar of field flowers, we simulated the flower environment to conduct several groups of behavioral experiments in the adult *Mythimna separata* by controlling the presence, absence or movement of pollens in nectar, then counted and compared the proboscis pollens.

We found that the pollens on the proboscis were mainly derived from the nectar.

Our conclusion may contributes to the research of pollen germination on stigmas, especially the dry stigma, and also shows the importance of adult Lepidoptera to pollination, even supports the coevolution of Lepidoptera and angiosperm.

## Introduction

Pollen germination is an important vital activity in plants. The germination of pollens needs sugar solution or nectar in nature as energy source (Broyles & Stoj, 2019; Hirsche et al., 2017; Portnoi & Horovitz, 1977; Mishal & Eisikowitch, 2022; Keven et al., 1989). Nectar is usually secreted by the nectary glands of the flower at the base of the petals, pistil, or stamens (Jennifer, 2009). Although some species’ flowers can secrete nectar on the stigma, but the stigma area is smaller than the area of nectar secreted by nectary. From this point of view, there can be rich pollens in the nectar, and this was supported by the filed data that pollens are ubiquitous in nectar investigated by Herrera (2017), including germinating pollens (Herrera, 2017; Sulborska-Różycka & Weryszko-Chmielewska, 2022). Consequently, it is basically impossible for pollen to touch the stigma when it has been inundated by nectar.

The role of adult Lepidoptera in plant pollination has been neglected or underestimated for a long time. Recent research has shown that these adults play important roles in pollination. In Mallorca, the major island of the Balearic archipelago in the western Mediterranean, researchers found that moths’ niche overlap was larger than that of the plants (Ribas-Marquès et al., 2022). In the Himalayan ecosystem of North-East India, settling moths were found to be the vital component of pollination (Singh et al., 2022). In the Yunnan Province of Southwest China, the researchers identified 554 species of pollinating butterflies, and their potential pollinating service could be a great contribution to montane agriculture, which has expanding areas of cash crops and fruit horticulture (Zhang et al., 2020). Except the above spatial research, the temporal research showed that the adult Lepidoptera were often complementary to the diurnal pollinators (Singh et al., 2022; Souza et al., 2022). The adult Lepidoptera even visit more flower species than bees (Walton et al., 2020). There is a general consensus that pollens are mainly deposited on the proboscis of adult Lepidoptera. The first record was done by Charles Darwin when he identified the pollinia of *Orchis pyramidalis* on the proboscises of nine Noctuidae specimens (Darwin, 1904). The research of other Lepidoptera, such as *Mamestra configurata, Heliothis zea, Agrotis ipsilon, Pseudaletia unipuncta* and *Mythimna separata* (Guo et al., 2018; Hendrix et al., 1987; Hendrix & Showers, 1992; Liu et al., 2017; Turnock et al., 1978), showed the same results, with all of their proboscises being the main deposition sites. The proboscis pollens can be transported to different plants for plant pollination (Arditti et al., 2012; Sugiura & Yamazaki, 2005). Based on this phenomenon, many researchers have used the identification of proboscis pollens to research the adult Lepidoptera’s nectar plants and their migrations (Courtney et al., 1982; Gregg et al., 2001; He et al., 2022; Hendrix & Showers, 1992; Kislev et al., 1972; Mikkola, 1971). However, it is not known how the pollens are deposited on the proboscis. Until now, there has only been a simple description that when adult Lepidoptera visit flowers, pollens are often involuntarily stuck to the proboscis (Jones & Jones, 2001; Jones, 2012), and the exact process remains unknown or overlooked.

There are two kinds of stigmas in plants, the wet stigma and the dry stigma (Broz & Bedinger, 2021; Doucet et al., 2016; Edlund et al., 2004). The wet stigma tends to trap and hydrate pollen indiscriminately, and pollens adhered to the wet stigma go through stages of hydration, germination, pollen tube penetrance of the stigma cuticle, and growth toward the transmitting tissue. (Broz & Bedinger, 2021). The germination of pollen on the dry stigma is complex, and only the *Arabidopsis thaliana* and several *Brassica* L. species have been researched. Pollen that is captured by a dry stigma depends on the biophysical or chemical properties of the pollen surface that promote adhesion, and the dry stigma accepts compatible pollen for fertilization, reject self-incompatible pollen to prevent inbreeding, and ignore foreign pollen (Broz & Bedinger, 2021; Edlund et al., 2004; Goring, 2018; Jamshed et al., 2020). The pollens that are adhered on the dry stigmas can’t acquire sugar and water for germination if they are rejected or ignored by the dry stigmas. However, almost all research has ignored the participation of insects (like the adult Lepidoptera) which may transport pollens from the nectar to stigmas, and that pollens may germinate on the dry stigma since they have soaked in the nectar. So, the key question is that whether insects carry pollens from nectar to the stigma.

By accident, we observed the sucking process and found a novel phenomenon that the pollens were mainly filtered outside and deposited on the proboscises of adult *M. separata* and *Helicoverpa armigera* (Video S1, S2). So, based on the above statements, the key question that whether insects carry pollens from nectar to the stigma can be verified by whether the pollens on the Lepidoptera proboscises are mainly derived from nectar.

Here, we tested the hypothesis that the pollens on the proboscis are mainly derived from nectar. In brief, we simulated flower environment by artificial flowers to conduct several groups of behavioral experiments on adult *M. separata* by controlling the presence, absence or movement of pollens in the nectar, then we compared the numbers of proboscis pollens among different groups and found that the pollens on the proboscis were mainly derived from the nectar.

## Material and Methods

### Insects

The *Mythimna separata* pupas were put in an incubator with a temperature of 25 ± 1 ℃, a humidity of 50 ± 10%, and photoperiod of 14:10 h light:dark (the photophase started at 05:00 AM and the scotophase started at 07:00 PM). Since the eclosion of *M. separata* pupas often happens at night, we removed the newly enclosed male adults at 05:00 AM and used them for experiments without feeding until the scotophase.

### Artificial flowers

The artificial flowers were made mainly referencing the methods of Good et al. (2014) and artificial anthers were added. In detail, red A4 paper was clipped into a wafer with eight petals, which was about 7 cm in diameter. The artificial anther was made as follows: pollen plates were prepared by pouring the rape pollen water solution onto filter papers and let them dry; a piece of copper wire about 10 cm long and 0.2 mm in diameter was cut; the wire was crimped to make both ends bigger untill it became 5 cm long; double-sided tape was wreathed on both ends and used as the anther bases, and it was about 1 cm long and 1.5 mm in diameter; and copper wire was rolled on a pollen plate to stick the pollens onto the anther bases. Subsequently, a 1.5 mL centrifuge tube cover was cut into a circle as a container for the nectar (the nectar solution was made up according to Nicolson [2007], and the constituents were 3% sucrose + 6% fructose + 6% glucose [w/v]). Finally, four copper wires were stuck onto the paper wafer using odorless hot glue to make one flower with eight artificial anthers, the cover was stuck in the middle to contain the nectar, and an iron wire was added as a pedicel.

### Feeding behavioral experiments to ensure the source of the deposited pollens on the proboscis

#### Basic operations of the behavioral experiments

All the experiments were conducted in the scotophase, at about 08:00-10:00 PM. One artificial flower was put in a can (10 cm in diameter, 10 cm in height) with a breathable cover. Then, 100 μL of the solution was added in the center cover, then one adult was put into the can for about 2 h. Finally, the moth was killed and the pollens on the proboscis were counted. All the artificial flowers were only used once.

#### The deposited pollens on the proboscises of the intact adults

Following the basic operations of behavioral experiments, one group was added nectar, and the other group was added rape pollen nectar solution (the pollen concentration was 2 μg/μL, which was similar to what Herrera observed in the field). Then moths were killed after about 2 h to count the pollen grains on the proboscis.

#### The deposited pollens on the proboscises of the wing-cut adults

The wings of adult *M. separata* were cut (the wings’ roots were preserved to prevent serious injury), then similar processes were conducted as above for the deposited pollens on the proboscises of the intact adults. Since the mobility of the wing-cut adults was impaired, the artificial flower pedicel was inserted into a foam platform to let the adults easily reach the nectar.

#### The deposited pollens on the proboscises of the intact adults that were feeding on non-free-flowing pollen nectar

Nectar was added in the center of the flower, then cotton fibers were put into the nectar to prevent the pollens which were blew off the anther from flowing freely. The basic operations of the behavioral experiments were followed, and the moths were killed after about 2 h to count the pollen grains on the proboscis.

#### The antenna pollens in the behavioral experiments

Since the antennae are the other parts that pollens adhere to in the adult moth (when compared with the pollens on the proboscis, the antenna pollens are lower in terms of the average number and percentage), the above five groups’ antenna pollens were also counted and compared to ensure whether the antenna pollens were related to the nectar pollens.

### Statistical analyses

All graphs were generated in Prism 6 (GraphPad Software, La Jolla, CA). T-test was used to analyze the statistical difference in SPSS 25 (SPSS Inc., Chicago, IL) (P<0.05).

## Results

The adults were found to suck nectar freely on the artificial flowers (Figure 1a). The result of the first feeding behavior experiment showed that the average amounts of proboscis pollens were no difference between the intact adults that consumed nectar and pollen nectar solution (Figure 1b). But when we examined the used flowers and the behaviors of the adult moths in the can, it was found that their flapping wings could stir air and blow the pollens off the anthers to pollute the nectar.

**Figure 1.**
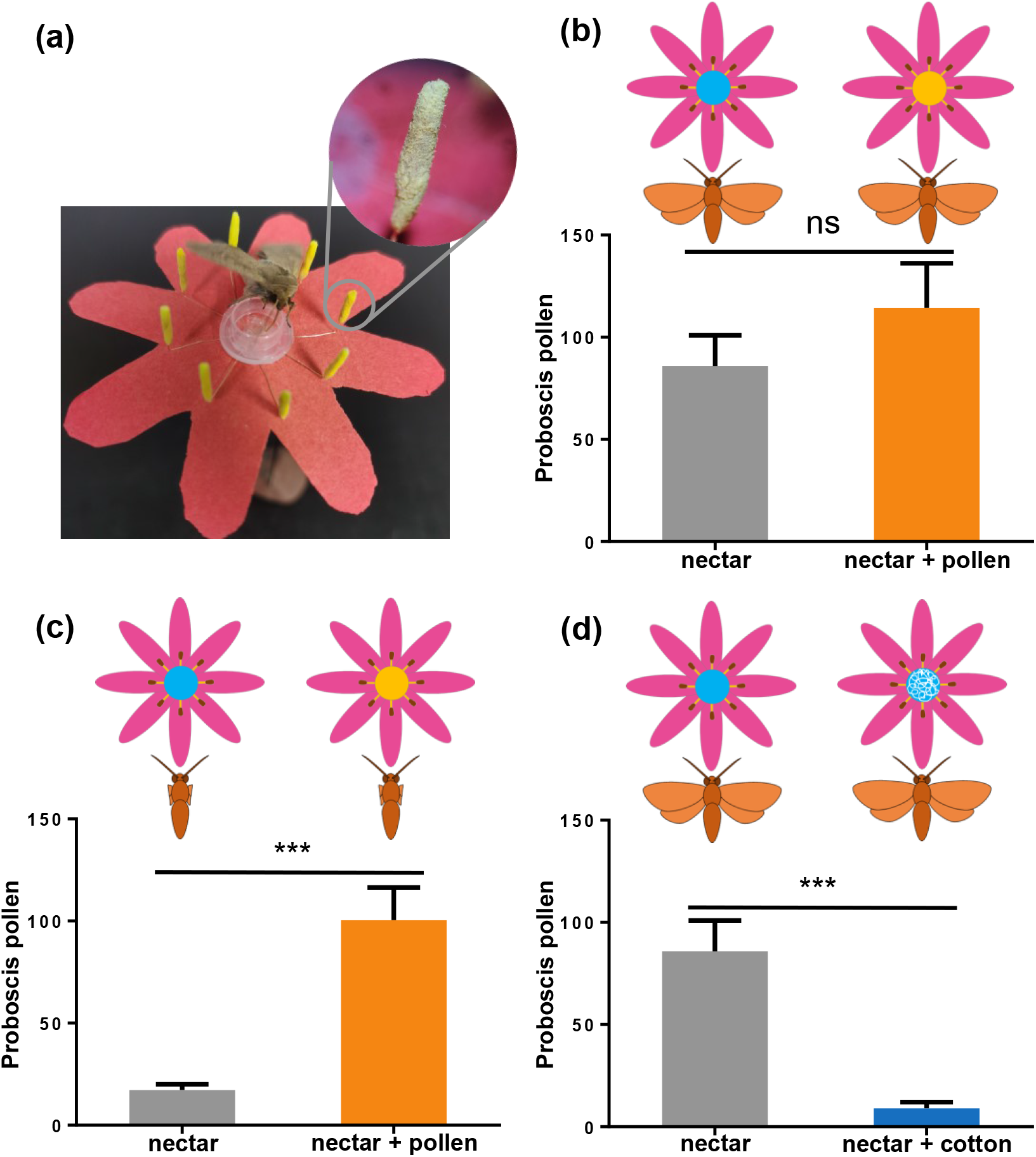
The average number of pollens on the proboscis for the different behavioral experiments. (a) The adult *Mythimna separata* sucking the nectar on the artificial flower, where the nectar is in the center centrifugal tube cover, the stamens are around the center, and the anther detail is showed in the circle. (b) The average amount of proboscis pollens between the intact adults that consumed nectar (gray, n = 34) and pollen nectar solution (orange, n = 34). (c) The average amount of proboscis pollens between the wing-cut adults that consumed nectar (gray, n = 30) and pollen nectar solution (orange, n = 30). (d) The average amount of proboscis pollens between the intact adults that consumed nectar (gray, n = 34) and cotton-added nectar (blue, n = 30). T-test was conducted for each pair, where the significance level was P<0.001.

Thus, the experiment was redesigned using two additional behavior experiments, where the first one prevented the anther pollens from being blown off by cutting the wings of the adults, and for the second one, cotton fibers were added into the nectar to prevent the pollens from flowing freely in the nectar. After the adult’s wings were cut, there were very few pollens that were deposited on the proboscises (Figure 1c). The adding of cotton fibers also prevented the pollens from flowing freely and very few pollens were deposited on the proboscises (Figure 1d).

There were five groups in the three kinds of behavior experiments (Figure 1), and when they were compared using t-test, the results showed that there were no differences in the number of the pollens on the proboscis between the intact adults and wing-cut adults that both consumed the pollen nectar solution (Figure S1).

Since the antennae are the other parts that pollens adhere to in adults, we also counted and compared each groups’ antenna pollens and found that the antennae pollens were fewer and not related to the nectar pollens (Figure S2).

## Discussion

The general consensus that pollens are mainly deposited on the proboscis of adult Lepidoptera has been viewed for more than a century, but the exact process of how the proboscis carries pollens has not been studied, even been overlooked. We have demonstrated that the proboscis pollens are mainly derived from nectar when the adult Lepidoptera sucks the nectar. Our finding shows that the insects (here are the adult Lepidoptera) can carry pollens from nectar to the stigma. It opens a new gate for the research of the pollen germination on the stigma, supports the coevolution of Lepidoptera and angiosperm, and also shows the importance of adult Lepidoptera in the pollination ecology and nectar ecology.

The germination of pollens needs sugar solution as energy source (Broyles & Stoj, 2019; Hirsche et al., 2017; Mishal & Eisikowitch, 2022), and the field investigation by Herrera (2017) showed that there were germinating pollens in the nectar of field flowers. So, the pollens which are rejected or ignored by the dry stigma may germinate if they are from the nectar, and our result “the Lepidoptera proboscis pollens are mainly derived from nectar” shows that the adult Lepidoptera can carry pollens from the nectar to the stigma. Thus, the botanists should consider this phenomenon in the further research of the pollen germination on stigmas, especially on the dry stigma.

In plants with dry stigmas, regulated pollen hydration provides an effective early barrier to incompatible pollination, and this mode is active in self-incompatible crosses and in crosses between species (Edlund et al., 2004). But it is not conducive to the evolution of angiosperm diversity, and in contradiction with the coevolution of Lepidoptera and angiosperm (Kawahara et al., 2019). However, if the pollens adhered on the dry stigmas are transported by the proboscises of adult Lepidoptera from the nectar, they may germinate directly, and this view can support the theory of the coevolution of Lepidoptera and angiosperm.

Overall, our result opens a gate for the botanists to research the pollen germination on the stigma, especially on dry stigmas, and supports the theory of the coevolution of Lepidoptera and angiosperm. It also promotes the understanding of pollination by adult Lepidoptera and the operating mechanism of pollination ecology and nectar ecology.

## Acknowledgements

This study was funded by National Natural Science Foundation of China (32130089), Shenzhen Science and Technology Program (Grant No. KQTD20180411143628272), Projects subsidized by Special Funds for Science Technology Innovation and Industrial Development of Shenzhen Dapeng New District (Grant No. PT202101-02), National Key R&D Program of China (2022YFE0116500), The Young Science and Technology Talent Support Project of Jilin Province (QT202121, QC). The authors would like to thank all the reviewers who participated in the review, as well as MJEditor (www.mjeditor.com) for providing English editing services during the preparation of this manuscript.

## Author contributions

BZR, GRW and BWM conceived and designed the research. BWM conducted the experiments. BWM, QC and FL analyzed the data. BWM, QC, BZR and GRW wrote and revised the manuscript. All authors read and approved the manuscript.

## Supplementary materials

**Figure S1 The comparative pollens on the proboscis among the five groups**. The average amount of proboscis pollens among the intact adults that consumed nectar (gray, n = 34), intact adults that consumed the pollen nectar solution (orange, n = 34), wing-cut adults that consumed nectar (blue, n = 30), wing-cut adults that consumed the pollen nectar solution (brown, n = 30), and intact adults that consumed the cotton-added nectar (green, n = 30). T-test was conducted for each two, where the significance level was P<0.001.

**Figure S2 The comparative pollens on the antenna among the five groups**. The average amount of pollens on the antenna among the intact adults that consumed nectar (gray, n = 34), intact adults that consumed the pollen nectar solution (orange, n = 34), wing-cut adults that consumed nectar (blue, n = 30), wing-cut adults that consumed the pollen nectar solution (brown, n = 30), intact adults that consumed cotton-added nectar (green, n = 30). T-test was conducted for each two, where the significance level was P<0.05.

**Video S1** The pollens were mainly filtered outside and deposited on the proboscis of an adult *Mythimna separata*.

**Video S2** The pollens were mainly filtered outside and deposited on the proboscis of an adult *Helicoverpa armigera*.

