## Supplementary figures and images for "Lepidoptera proboscis pollens are mainly derived from nectar"

### Supplemental Figure 1

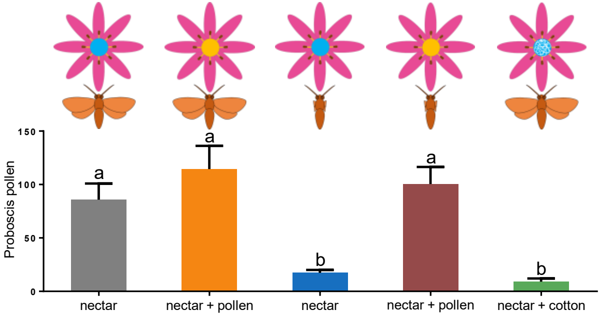

### Supplemental Figure 2

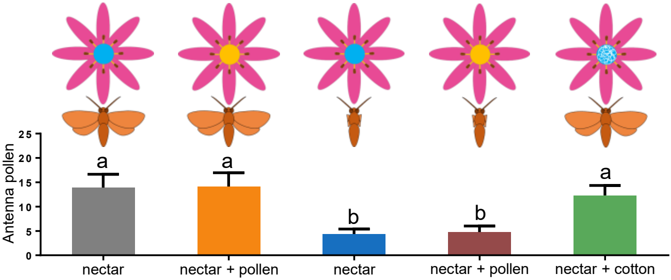
